# GerAB residues predicted to interfere with water passage based on steered Molecular Dynamics are key to germinosome stability

**DOI:** 10.1101/2024.06.28.601167

**Authors:** Longjiao Chen, Houdijn Beekhuis, Christina van den Bosch, Gianni Vinay, George Korza, Jocelyne Vreede, Peter Setlow, Stanley Brul

## Abstract

Some *Bacillales* and *Clostridiales* bacteria form spores in unfavorable environments. These spores are dormant but can rapidly resume metabolism in response to certain small molecule nutrients. This process is termed germination and can be initiated by a variety of low molecular wt nutrients termed germinants. Structural modeling and mutagenesis studies showed that GerAB, an inner membrane (IM) protein of the *Bacillus subtilis* spore germinant receptor (GR) GerA, is involved in L-alanine-initiated spore germination. A previous molecular simulation study also suggested there is a water channel in GerAB. In the current work, Steered Molecular Dynamics (SMD) simulations were employed to force a single water molecule through GerAB, identifying three key amino acid residues, Y97, L199 and F342, that interfere with water passage. When these residues were altered to alanine, L-alanine germination was minimal in spores with L199A, F342A and triA (Y97A, L199A and F342A triple mutant), while Y97A mutant spores germinated ∼61%. And besides Y97A, all other mutants showed compromised germination triggered by AGFK mixture. Western blotting found reduced levels of the GerA GR in the Y97A mutant, and an absence of the GerA GR in all other mutants. This proved that all three identified residues are crucial to the structural integrity of the GerAB protein and therefore probably the formation of the GR complex, the germinosome.

## Introduction

Several species within the *Bacillales* and *Clostridiales* bacterial orders can form dormant spores in harsh environments. These spores are metabolically inert, extremely resistant to a variety of harsh treatments, and dormancy can last for decades. However, cells derived from some of these spores can cause food spoilage, infectious diseases, intoxications and foodborne illnesses^1^. This is especially true for spores of *Bacillus anthracis, Clostridiodes difficile* and *Bacillus cereus*. Consequently, the biology of spore-forming bacteria has been investigated for decades. A major feature of dormant spores is their low core water content, 25%-45% of wet weight in spores of the model organism *Bacillus subtilis*, compared to ∼80% of wet weight in its growing cells. However, how water is taken up into the dormant spore core is not clear, as *B. subtilis* lacks known water pores such as aquaporins^2,3^.

The loss of spore dormancy and resumption of its metabolism and cell-like structure requires spore germination. Small molecule nutrients, termed germinants, including specific monosaccharides, monovalent cations, and amino acids, can trigger spore germination with no germinant catabolism involved^4^. Germinant receptors (GRs) play a crucial role in kick-starting germination. There are three GRs in the model organism *B. subtilis*, GerA, GerB, and GerK. Each GR is composed of three subunits, A, B, and C, and requires the presence of all three subunit to function^1,4^. GRs and GerD protein colocalize into cluster termed germinosome in the spore inner membrane (IM). Even though its detailed structure is unknown, its structural stability is necessary for rapid and cooperative response of nutrients through GRs^1^. The B subunit of GerA, GerAB, is a transmembrane protein belonging to the APC transporter superfamily that mediates L-alanine-triggered germination^5–7^. Similar function and organization were identified for GerB and GerK except these two collectively respond to a mixture of AGFK (L-asparagine, D-glucose, D-fructose and K^+^ ions)^13^ instead. Many attempts have been made to fully understand the structure and function of GRs. Recently, GerAB has been shown to be the ligand sensing subunit ^7^ using structure prediction and mutagenesis. With Molecular dynamics (MD) simulations, Blinker *et al*. observed water passing through GerAB, thus identifying a putative water channel ^8^. A recent study by Gao *et al.* combined AlphaFold^9^ structure prediction and mutagenesis, showing that GerA forms an oligomeric membrane channel which releases monovalent cations upon sensing L-alanine^10^. However, to date, the function and mechanism of the putative water channel in GerAB and its role in spore water intake during germination not been fully elucidated and experimentally verified.

To further investigate the molecular mechanism of the putative water channel in GerAB, MD simulation studies coupled with experimental mutagenesis and spore germination assays have been carried out in the current work. Water molecules were pulled through GerAB from the outside of the spore IM to its inside with Steered Molecular Dynamics (SMD) simulations. In these simulations, the termini of GerAB are assumed to be located on the inside of the spore^10^ (Figure 1). During SMD simulations, the work required for pulling a water molecule through GerAB was measured. Visual inspection of the SMD trajectories showed three amino acid residues Y97, L199 and F342 interfere with the passage of water through GerAB. The sidechains of these residues are close to one another and act as obstacles to water passage. To verify the role of these residues, spores of mutants Y97A, L199A or F342A and the triA (Y97A, L199A and F342A triple mutant) were constructed, and the germination efficiencies of the wild type (wt) and mutant spores were measured. With L-alanine as the germinant, wt spores germinated close to 100% and Y97A mutant spores germinated around 61%, while L199A, F342A and triA mutant spores germinated less than 1%. When using AGFK as the germinant, all mutant spores germinated at comparable level to wt spores, with at most an ∼10% germination efficiency drop. Western blotting verified both wild type and Y97A spores contained GerAA, with Y97A mutant spores showed a reduced level. However, the other three mutant spores completely lacked GerAA. This indicates that only the wild type and Y97A mutant spores could assemble the complete GerA GR, albeit Y97A mutant spores to a lower amount. These observations collectively demonstrate that the three identified residues, Y97, L199, and F342 are important in the structural integrity of GerAB, therefore the assembly of the GerA GR, and thus the stability of the IM GR complex germinosome.

**Figure 1.**
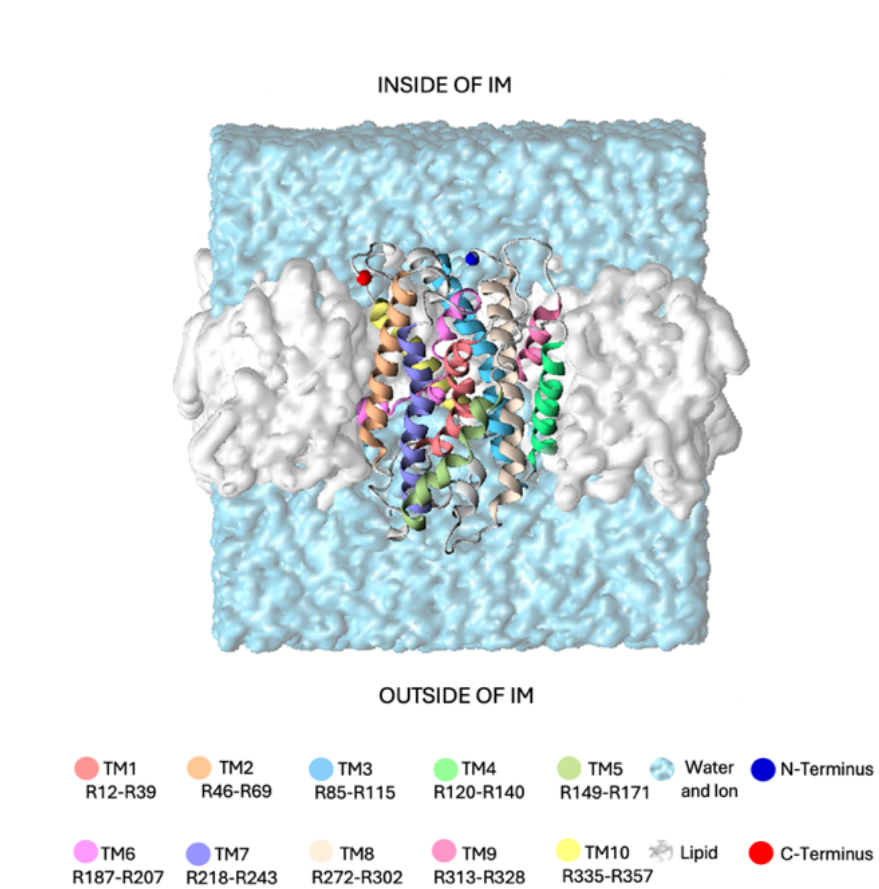
Simulation system used in this study. The GerAB protein is embedded in a lipid bilayer membrane (shown as a white surface), with its N-and C-termini oriented towards the interior of the spore’s inner membrane (IM). The protein is displayed as a ribbon, with the 10 transmembrane regions (TMs) individually color-coded. Residues assigned to each TM follow the prediction by Blinker *et al*.^8^ The surrounding water, containing 20 mM KCl, is represented as a light blue surface.

## Results

### 1. Key residue identification

In the SMD simulations, water molecules were pulled through GerAB from the outside of the spore IM to the inside of the spore IM, as indicated in Figure 2A. For each individual SMD run, the cumulative work as a function of the z coordinate was calculated. For all runs, the exerted work increases at z = 5 nm, until z = 7.5 nm. This means that the water molecule does not spontaneously go through the protein at the pulling speed. At z > 7.5 nm, the cumulative works stays approximately constant, which means that no additional work is required to pull the water molecule through bulk water after it has exited the protein (Figure 2B). This observation shows that the SMD does not suffer from artifacts caused by pulling the water too fast, as pulling the water molecule through bulk water happens at a velocity typical for water diffusion at 298 K^11^. This indicates that the pulling speed is sufficiently slow.

**Figure 2.**
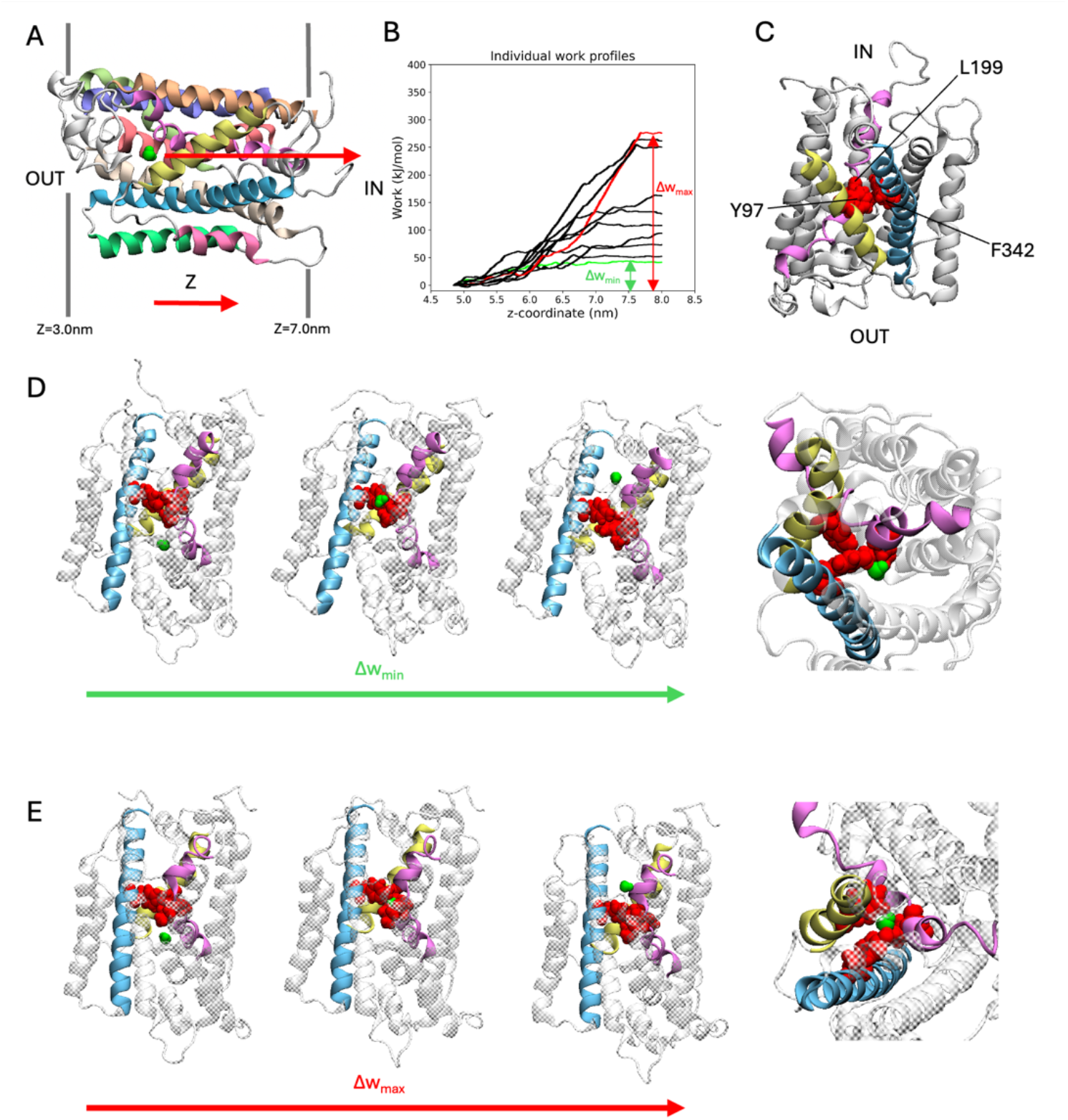
SMD Simulations of water molecules pulled through GerAB. (A) Example of the starting frame for the SMD simulations. The water molecule which is pulled is shown in a green space filling representation. The arrow indicates the pulling direction, from the outside of the spore IM to the inside. Dark grey lines flanking the protein indicate the membrane. (B) Work profiles of the SMD simulations along the z-axis. The work profiles Δw_max_ and Δw_min_ are shown in red and green, respectively. (C) A triad of Tyr97, Leu199, and Phe342 (all in red space filling representation) are located on TM3, TM6, and TM10, respectively. Only TMs with the triad are colored, according to the coloring scheme in Figure 1. Protein regions not part of the triad is shown in white. (D), (E), Water molecule passing the triad in the SMD simulation with Δw_min_ and Δw_max_, respectively. From left to right: snapshots of simulation for water oxygen at z = 4.84 nm, 5.89 nm, 6.65 nm (for Δw_min_), and z = 5.07 nm, 6.21 nm, 7.84 nm (for Δw_max_), showing the water before, during, and after passing the triad, followed by a closeup of the triad with the water molecule during the water molecule passing the triad. The closeup is shown from the inside of the spore.

The pulling work profiles all have a similar shape but differ in when the plateau is reached. To quantify this variation, the difference of the cumulative work is defined as Δw = w_end_ – w_start_. Δw ranged from Δw_min_= 42 kJ/mol to Δw_max_= 275 kJ/mol (Figure 2B). According to the Boltzmann distribution, the process requiring the least amount of work is the most likely to occur during the process of water passage^12^. Consequently, we visually inspected the run with the smallest Δw, labeled Δw_min_. In this simulation, a group of three amino acids with bulky side chains (termed the triad, consisting of Tyr97, Leu199, and Phe342, Figure 2C) blocked the movement of the water molecule along the z-axis. To continue moving through GerAB, the water molecule bypassed the triad between TM3 and TM6 where Tyr97 and Leu199 are located, respectively (Figure 2D). In contrast, in all the other SMD simulation trajectories, the water was pulled through the three bulky side chains of the triad, with run Δw_max_ as an example (Figure 2E).

In summary, the bulky side chain triad interfered with water molecule passage in all the SMD simulations trajectories, albeit with different water paths. To verify the function of the triad *in vivo*, the triad residues have been mutated to alanine to test the function of the bulky side chains on germination behavior of *B. subtilis* spores.

### 2. Y97, L199 and F342 are important to the structural stability of germinosome

Four mutant strains of *B. subtilis* PY79 were successfully constructed, including Y97A, L199A, F342A, and triA (Y97A, L199A and F342A triple mutant). Differences in germination efficiency of each strain with L-alanine or AGFK shows that each residue in triads has an important role in both L-alanine mediated germination and AGFK initiated germination due to the reduced ability of mutant spored in respond to both types of germinants compared to wt spores. In the presence of L-alanine, only Y97A mutant spores are viable and capable of germinating partially with a germination efficiency of 61%, while the other mutants failed to respond to L-alanine (Figure 3A). Similar pattern is shown in OD600 drop where only Y97A spores showed partial OD drop while other mutants exhibited none (Figure 4A). All mutants respond to AGFK in a slightly lagged kinetics compared to wt, with an insufficient OD_600_ drop indicating a germination efficiency drop, albeit with different level in different mutant spores (Figure 4B). This is in consensus with single-spore germination efficiency assay, which revealed a drop around 10% of germination efficiency in L199A, F342A and triA mutant spores (Figure 4A). Notably, the insufficiency of OD_600_ drop and the germination efficiency drop in Y97A are both less than that of the other mutants.

**Figure 3.**
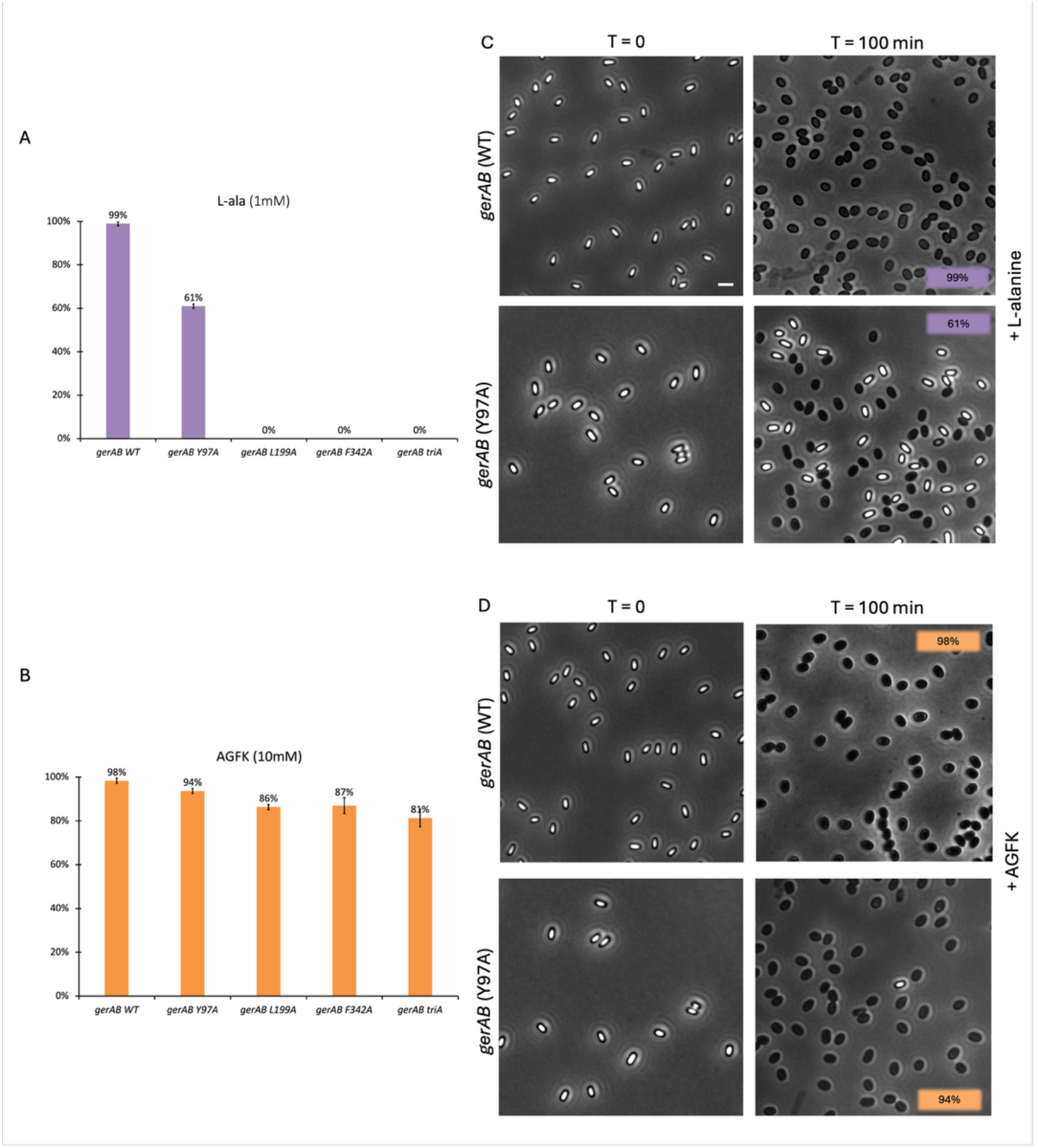
Single spore germination assay of wt and different spore variants by phase contrast microscopy. Germination efficiency of spores with different GerAB variants with L-alanine (A) or AGFK (B). Wild type spores’ germination was close to 100% with both germinants. In responds to L-alanine, only Y97A germinates 61% while other mutants are unable to germinate. In respond to AGFK, Y97A spores exhibits the highest spore viability among all four mutants. (C), (D) Representative phase-contrast microscopy images of wt and Y97A spores before and after a 100 min incubation with L-alanine or AGFK, respectively. Dormant spores are phase bright while germinates spores are phase dark under the current microscopy condition. Scale bar, 2 μm. Error bars in A and B indicate ± SD of three technical replicates. Uncropped microscopy images are presented in Figure S1.

**Figure 4.**
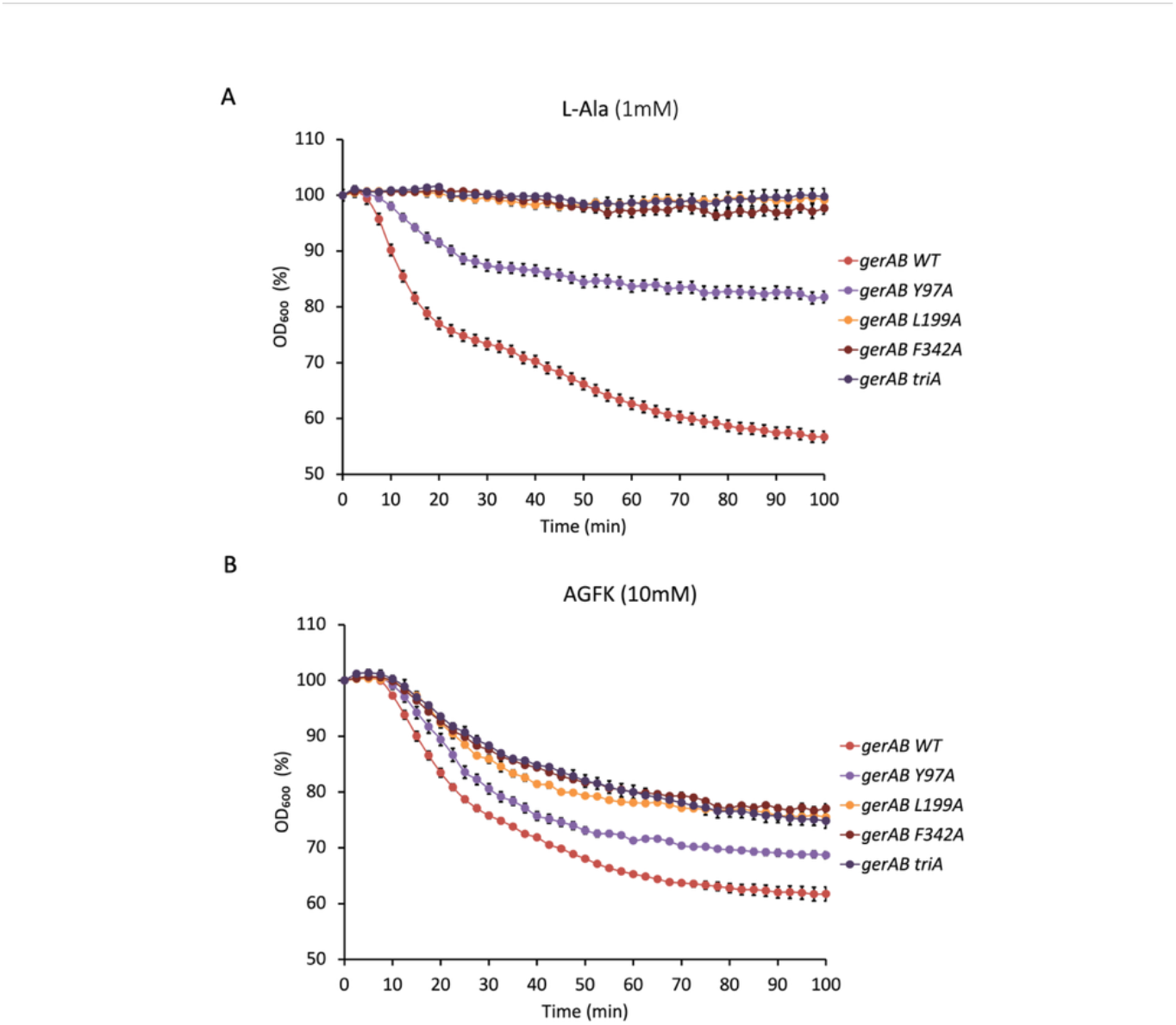
Spore germination assay with OD_600_ drop with 1mM L-alanine (A) or 10mM AGFK (B). Y97A mutant spores showed partial germination compared to wt spores in response to L-alanine. L199A, F342A and triA mutant spores failed to germinate in the presence of L-alanine. All mutants showed compromised germination with AGFK, although less so in Y97A mutant. Error bars indicate ± SD of three technical replicates.

In addition to analysing spore germination, we also performed Western blot analysis of spore proteins with a purified GerAA antibody to assess the amount of GerA GR in spores of each variant. Only wt and Y97A mutant spores exhibited GerAA signals, although the level in the Y97A mutant is lower than that in wt spores (Figure 5). Since assembly of the GerA GR requires all three subunits, this means that the low response to L-alanine of L199A, F342A and triA mutant spores was due to the absence of whole GerA GR^15^. That is to say GerAB Y97A is produced and assembles into a complex with GerAA and GerAC, albiet the lower level indicates its lower stability. At the same time, GerAB L199A, GerAB F342A and GerAB triA are either unstable or unable to form a complex with GerAA and GerAC proteins. This indicates all three amino acid sites are important to the structural stability of GerAB. And compared to Y97, L199 and F342 are even more important structurally in germinosome formation, since their mutagenesis completely eliminated the existence of the GerA GR.

**Figure 5.**
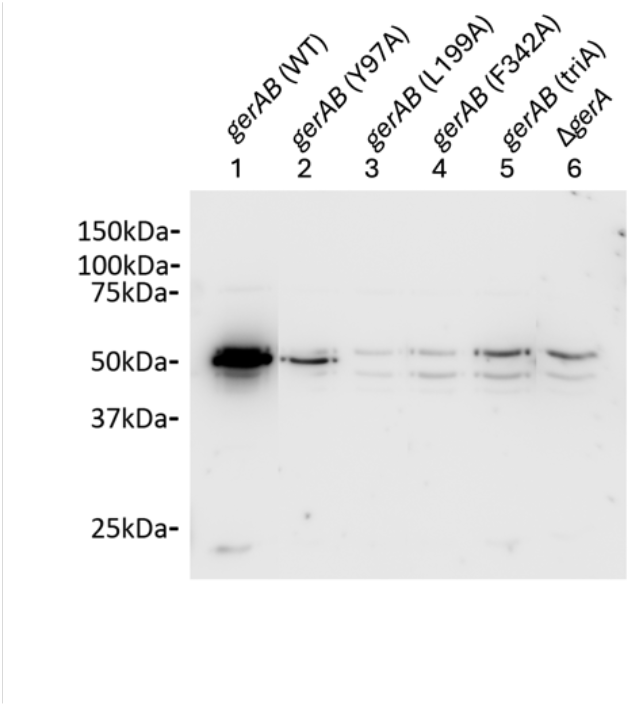
Western blot of *B. subtilis* PY79 wt and mutant spore proteins with anti-GerAA antibody. Both *gerAB* (wt) and *gerAB* (Y97A) sample show a clear band with the GerAA antibody at ∼50 kDa, while the other mutants showed no GerAA band. However, *gerAB* (L199A), *gerAB* (F342A) and *gerAB* (triA) spores exhibited only the same bands as the Δ*gerA* GerAA deletion control sample. All samples were taken from the same blot; irrelevant lanes were removed. Uncropped blot image is presented in Figure S3.

Additionally, only spores of the Y97A mutant germinated with AGFK identically to that of wt spores in single spore germination efficiency assay (Figure 3C), and Y97A mutant spores exhibited germination kinetics against AGFK the most similar to that of wt (Figure 4B). This is surprising since AGFK is sensed by the GerB and GerK, GRs with no direct involvement of GerAB. Consistent with the GerA GR only present in the Y97A mutant, this indicated that the absence of the GerA GR could affect the stable assemble of gernimosome sicne the ability of GerB and GerK GRs respond to AGFK were also compromised in L199A, F342A and triA mutant. Ultimately, the deletion of *gerA* operon also led to a compromised germination in respond to AGFK since the there is a drop of DPA release of *ΔgerA* strain compared to that of WT strain (Figure S2). These observations are consistent with the current understanding of germinosome function since different GRs in *B. subtilis* spores form protein complex in germinosome and work synergistically ^16,17^. The current study thus provided evidence that the triad is also structurally important in the assembly of the whole germinosome, not only the GerA GR.

## Discussion

In this study, the function of GerAB was studied by both MD simulations and mutagenesis. By pulling a single water molecule through GerAB in SMD simulations, we discovered that the side chains of Y97, L199 and F342, termed the triad in this study, interfere with water passage. The SMD further revealed two different ways of interaction between water molecule and the triad. In the simulation with the lowest pulling work (Δw_min_= 42 kJ/mol), the water molecule moved through GerAB bypassing the triad between TM3 and TM6. The simulation with the highest pulling work (Δw_max_= 275 kJ/mol) showed the water molecule passing through the three bulky residue sidechains of the triad. The qualitative observations of simulation in this study pointed out the role of triad residues in blocking water passing through GerAB, and indicated there may be different mechanisms for water passage through GerAB. For better quantitative results, *i.e.* a free energy profile, more SMD simulation runs need to be carried out, to reduce variance.

On the other side of the work, using mutagenesis of GerAB followed by germination assays and western blot analysis, we verified that Y97, L199 and F342 are crucial to the structural integrity of GerAB. In the Y97A mutant, both the efficiency of L-alanine mediated germination and the level of GerA GR are lower than that in the wt spores, while the other mutations eliminated stable GerA GR formation. The lack of the GerA GR in L199A, F342A and triA spores also led to a lower AGFK germination efficiency. This result suggests that our predicted residues in GerAB are very important structurally in germinosome formation. However, what this result did not confirm is our hypothesis that the triad is interfering with water passage. Unfortunately, since all four mutants failed to form a GerA complex at the level in wt spores, it is not possible to verify the role of the triad in L-alanine-initiated germination, not to mention the predicted new function, namely the water channel formation.

To further prove the exact role of triads in the function of GerAB, mutating the three residues to different residues (other than alanine) might give us further insight, since other mutants might form a stable GerA complex. However notably, investigation of the possibly water passage of GerAB as a way of water intake during spore germination is out of the scope of current study, since it is almost impossible to directly prove experimentally how and if water transportation happens through GerAB. With the combination of simulation, future work could provide more detailed and quantitative understanding of the mechanism of the interaction between water and GerAB, and how these interactions contribute to other molecular events during L-alanine-initiated spore germination.

## Materials and Methods

### 1. Simulations and analysis

One 100 ns molecular dynamics (MD) simulation of a protein structure model based on *Methanocaldococcus jannaschii* ApcT (PDB: 3GI8)^19^ generated by the structure prediction tool raptorX^20^ was conducted using the same starting structure and settings as those described by Blinker *et al*^8^. The simulation was performed with GROMACS 2020.4^21^, using the CHARMM36^22^ forcefield. In a cubic periodic box with starting dimensions of 10 by 10 by 11 nm, the system was solvated by TIP3P water molecules^23^. The simulation was maintained at a temperature of 298 K using the Bussi velocity rescaling thermostat^24^ and at constant pressure of 1.0 bar with the Parrinello-Rahman barostat^25,26^. Long-range electrostatic interactions were treated using the Particle Mesh Edward^27^ method with a maximum grid spacing at 1.6 Å. For short-range electrostatic and Van-der-Waals interactions, the cutoff radius was set to 12 Å. The production run lasted 100 ns with a timestep of 2 fs, with coordinates saved every 2 ps. The term “(free) MD simulation” refers to the MD simulation production run conducted for this research.

Steered Molecular Dynamics (SMD) simulations were performed to pull a water molecule through the GerAB protein. To extract suitable starting structure from MD simulation trajectory, the protein was first positioned at the center of the simulation box (with its center of mass between 4.76 nm and 4.85 nm along the x-and y-coordinate, and membrane boundaries at 3.0 nm < z < 7.0 nm). The entrance of the water was defined within the region 3.5 nm < x < 6.5 nm, 3.5 nm < y < 6.5 nm, and 4.6 nm < z < 5.0 nm. Ten frames, each containing a single water molecule within the defined entrance region, were extracted as the starting configurations for steered molecular dynamics (SMD) simulations. The selected water molecule was then subjected to the pulling force during the simulation. The steered MD simulations were conducted using PLUMED 2.7.0^28–30^ embedded with GROMACS 2020.4, where the steering collective variable (CV) was the position in the z-direction of the oxygen atom of the pulled water molecule. In all SMD simulations, the selected water molecule was pulled 4 nm over 100 ns along the z-coordinate towards the inside of the inner membrane (IM), following the direction of water intake during germination. Over the 100 ns simulation, a moving harmonic potential kept the water oxygen constrained to z with a force constant κ = 10000 kJ mol^-1^ nm^-2^. The harmonic restraint moved at a constant velocity of 0.04 nm/ns. In addition, the water oxygen atom was constrained in the x and y direction with a harmonic potential with a force constant κ = 10 kJ mol^-1^ nm^-2^, to keep the water molecule inside the protein. The SMD simulations were performed with the same settings as the MD simulation. The work performed to constrain the water molecule along the z coordinate in the SMD simulations was computed as the cumulative force applied to the water molecule over the simulation time. In total, 10 SMD runs were performed. Visual inspection of the simulation trajectories was conducted using Visual Molecular Dynamics (VMD) 1.9.4a53 version^31^.

### 2. Mutagenesis

All strains were derived from *Bacillus subtilis* PY79. The *gerA* operon was cloned by PCR, inserted in plasmid vector pUC19 with the Gibson Assembly Master Mix kit (New England BioLabs, NEB # E2611S). Mutagenesis was carried out with the QuikChange Lightning Site-Directed Mutagenesis Kit (Agilent Technologies, Cat # 210518-5). Mutant *gerAB* sequences were integrated into the original *gerAB* locus by double crossover along with an erythromycin (erm) resistance cassette. The precision of the mutagenesis in the *gerAB* locus was confirmed by Sanger sequencing of all mutants. Strains involved in this study are listed in Table 1.

**Table 1.** Strains used in this study.

| Strain | Strain | Source |
| --- | --- | --- |
| *gerAB* Wild Type (wt) | *Bacillus subtilis* PY79 | Lab stock |
| *gerAB* Y97A::erm | *Bacillus subtilis* PY79 | Constructed in this study |
| *gerAB* L199A::erm | *Bacillus subtilis* PY79 | Constructed in this study |
| *gerAB* F342A::erm | *Bacillus subtilis* PY79 | Constructed in this study |
| *gerAB* triA::erm | *Bacillus subtilis* PY79 | Constructed in this study |
| $\Delta gerA$ | *Bacillus subtilis* PS832 <sup>35</sup> | Prof. Peter Setlow lab, UConn Health |
| $\Delta gerA$::spec | *Bacillus subtilis* PS832 <sup>35</sup> | Prof. Peter Setlow lab, UConn Health |

### 3. Sporulation and spore purification

Bacterial strains were streaked onto LB agar plates with appropriate antibiotics and incubated overnight to avoid stationary phase; a single colony was inoculated into liquid LB with antibiotics. Once OD_600_ reached 1.0-2.0, 200 µL liquid culture was spread onto 2xSG agar plates (Difco Nutrient Broth 16 g/L, KCl 26 mM, MgSO_4_ 2 mM, MnCl_2_ 0.1 mM, FeSO_4_ 1.08 µM, Ca (NO_3_)_2_ 1 mM, Glucose 5.5 mM, Agar 15g/L) without antibiotics. Plates were incubated upside down in plastic bags at 37°C for 2 to 5 days to allow sporulation, monitored by phase-contrast microscopy. After sporulation, the plates were dried on the benchtop for approximately 2 days to allow cell lysis. Spores were scraped from the plates and transferred to 50 mL centrifuge tubes with cold MQ water. The spore suspensions were then sonicated for 1 minute at full power, cooled on ice, and centrifuged at ∼8,000 rpm for 20 minutes to remove debris. The purity of spore preparation was checked with phase contrast microscopy.

### 4. Germination assay with phase contrast microscopy

Purified phase-bright spores at OD_600_ of 5 in MQ water were heat-activated at 70°C for 30 min followed by 20 min incubation on ice. Before the addition of the germinants, the germinant solutions were kept on ice. A final concentration of 1 mM L-alanine or 10 mM AGFK solution (10 mM each of L-asparagine, D-glucose, D-fructose and KCl)^7,13^ was added prior to slide preparation and imaging according to Wen et al.^32^ In essence, to prepare a slide for wide field imaging, slides and two types of coverslips (size 22mm round and size 22mm by 30mm rectangular) were cleaned with 70% EtOH and air dried in a vertical position minimizing dust collection. Two rectangular coverslips were prewarmed for several seconds on a 70°C heating block, 65 μl 2% agar was deposited on top of one coverslip, while the other coverslip was placed on top of it, spreading the agar in between. The agar patch was dried for approximately 10 min, after which the coverslips were slid off each other. The agar patch was cut into a 1×1cm section to which 0.4 μl of purified spore suspension with germinant added. The patch was transferred onto a round coverslip by placing it onto the patch and sliding it off. A G-frame was stuck onto the air-dried slide, onto which the coverslip was placed, closing all corners of the frame, and completing the slide for microscopy. For wide field time-lapse experiments, we used a Nikon Eclipse Ti equipped with a Nikon Ti Ph3 phase contrast condenser. Connected to it were a Nikon Plan Apo Plan Apo λ Oil Ph3 DM lens (100X, NA=1.49, T = 23°C), a Lumencor Spectra Multilaser (470 nm, 555 nm) using Lambda 10-B filter blocks, a NIDAQ Lumencor shutter, a Ti XY-and Z drive, and a Hamamatsu C11440-22C camera. All hardware was connected to a computer running NIS-Elements AR 4.50.00 (Build 1117) Patch 03. Before every use the condenser was adjusted to the optimal setting bringing the pixel size to 0.07 μm. A 100-minute time lapse video was recorded for every slide with a 30 sec interval between frames. The slide chamber was kept at 37°C, and single-cell germination was analyzed with SporeTrackerX^33^. The percentages of germinated spores at the end of the time-lapsed session were defined as germination efficiency.

### 5. Germination assay with optical density

Purified phase-bright spores, normalized to an OD_600_ of 1.2 in 25 mM HEPES buffer (pH 7.4), were heat-activated at 70 °C for 30 minutes, followed by incubation on ice for 20 minutes. 100 μL of the heat-activated spores were added, and germinant solutions were dispensed into a 96-well plate to a final concentration of 1mM for L-alanine and 10mM for AGFK. The OD_600_ was monitored every 2.5 minutes for 100 minutes using a plate reader, with the plate maintained at 37°C and agitated between measurements. The percentage of OD drop is measured against the initial OD. Every experiment was repeated 3 times and the average OD drop for each condition is shown.

### 6. Germination assay with DPA release

CaDPA release was measured using a fluorescence plate reader following the method outlined by Yi and Setlow. ^5^ Spores, at an OD_600_ of 0.5, were germinated with 1 mM L-alanine in 25 mM K-HEPES buffer (pH 7.4) at 45°C without prior heat activation. At various time points, 190 μL of the sample was mixed with 10 μL of 1 mM TbCl_3_, and relative fluorescent units (RFU) were recorded using a plate reader.

### 7. SDS-PAGE and immunoblotting

To prepare samples for western blotting, the spores were first decoated and then lysed as follows. 50 ODs of spores were pelleted by centrifugation. The pelleted spores were resuspended in 1 mL TUDSE buffer (8 M Urea, 50 mM Tris-HCl, pH 8.0, 1% SDS, 50 mM DTT, 10 mM EDTA) and incubated for 45 min at 37°C, centrifuged (3 min, max rpm, room temperature) and resuspended in 1 mL TUDS (8 M Urea, 50 mM Tris-HCl, pH 8.0, 1% SDS) buffer. The suspension was then incubated for another 45 min at 37°C, centrifuged (3 min, max rpm, room temperature) and then washed six times by centrifugation (3 min, max rpm, room temperature) and then all suspended in 1 mL TEN buffer (10 mM Tris-HCl pH 8, 10 mM EDTA, 150 mM NaCl). The final decoated spores were resuspended in 1mL water if samples were not immediately given further treatment. 50 OD of the decoated spore preparation was treated with 1mg lysozyme in 0.5mL TEP buffer (50 mM Tris-HCl pH 7.4, 5 mM EDTA) containing 1mM phenylmethylsulfonyl fluoride (PMSF), 1μg RNase, 1 μg DNase I, and 20 μg of MgCl_2_ at 37°C for 6 to 8 min, and then put on ice for 20 min. Glass disruptor beads (0.10 to 0.18 mm, 100 mg) were added to each sample and spores were disrupted with three bursts of sonication (micro probe, medium power for 10 sec, and placed on ice for 30 sec between bursts). Following the final sonication, samples were allowed to settle for 15 sec and 100 μl of the upper liquid was withdrawn and added to 100 μl 2x Laemmli sample buffer (BIO-RAD cat. #1610737) containing 5% (v/v) 2-mercaptoethanol and 1 mM MgCl_2_ and incubated at 90-95°C for 3 min. This was saved as the total lysate and ready to run on SDS-PAGE. 20 μg of total protein was loaded onto each lane of SDS-PAGE on a 10% acrylamide gel (BIO-RAD cat. #4561034) and run 45 min at 60 V followed by 45 min at 110 V. After SDS-PAGE, proteins on the gel were transferred to a 0.22 μm PVDF membrane. The membrane was then blocked for 30 min by 2.5% low fat milk in TBST buffer (20 mM Tris, 150 mM NaCl, 0.1% Tween 20) and incubated with 1:3000 anti-GerAA antibody overnight^34^. After washing three times with TBST, the membrane was incubated with 1:2500 diluted goat anti rabbit HRP antibody (BIO-RAD cat. #1706515) for an hour before visualization.

## Acknowledgements

I would like to thank my supervisors, Jocelyne, Peter and Stanley for their invaluable guidance and support throughout this project. I would also like to acknowledge the China Scholarship Council (CSC File No. 202109120009) for providing the funding that made this work possible.

## Author contributions statement

L.C. conducted simulations and analysis, conducted experiments on sporulation and germination assays, and wrote the manuscript. H.B. conducted simulations and analysis. C.B. carried out mutagenesis experiments. G.V. performed sporulation and germination assays. G.K. performed Western Blot analyses. J.V. provided supervision and guidance on the simulations. P.S. and S.B. provided supervision and oversight on spore biology. J.V., P.S. and S.B. contributed to the study design, and secured funding, and critical revisions and feedback on the manuscript.

## Data availability statement

Input files for the SMD simulations reported here are deposited on FigShare with https://doi.org/10.6084/m9.figshare.27225249.v1. Representative large-field time-lapse videos are deposited on FigShare with https://doi.org/10.6084/m9.figshare.27226959.v1.

