## Supplementary material for "GerAB residues predicted to interfere with water passage based on steered Molecular Dynamics are key to germinosome stability": Figure S1; Figure S2; Figure S3

**Supplementary information**

Input files for the SMD simulations reported here are deposited on FigShare with https://doi.org/10.6084/m9.figshare.27225249.v1. Representative large-field time-lapse videos are deposited on FigShare with https://doi.org/10.6084/m9.figshare.27226959.v1.

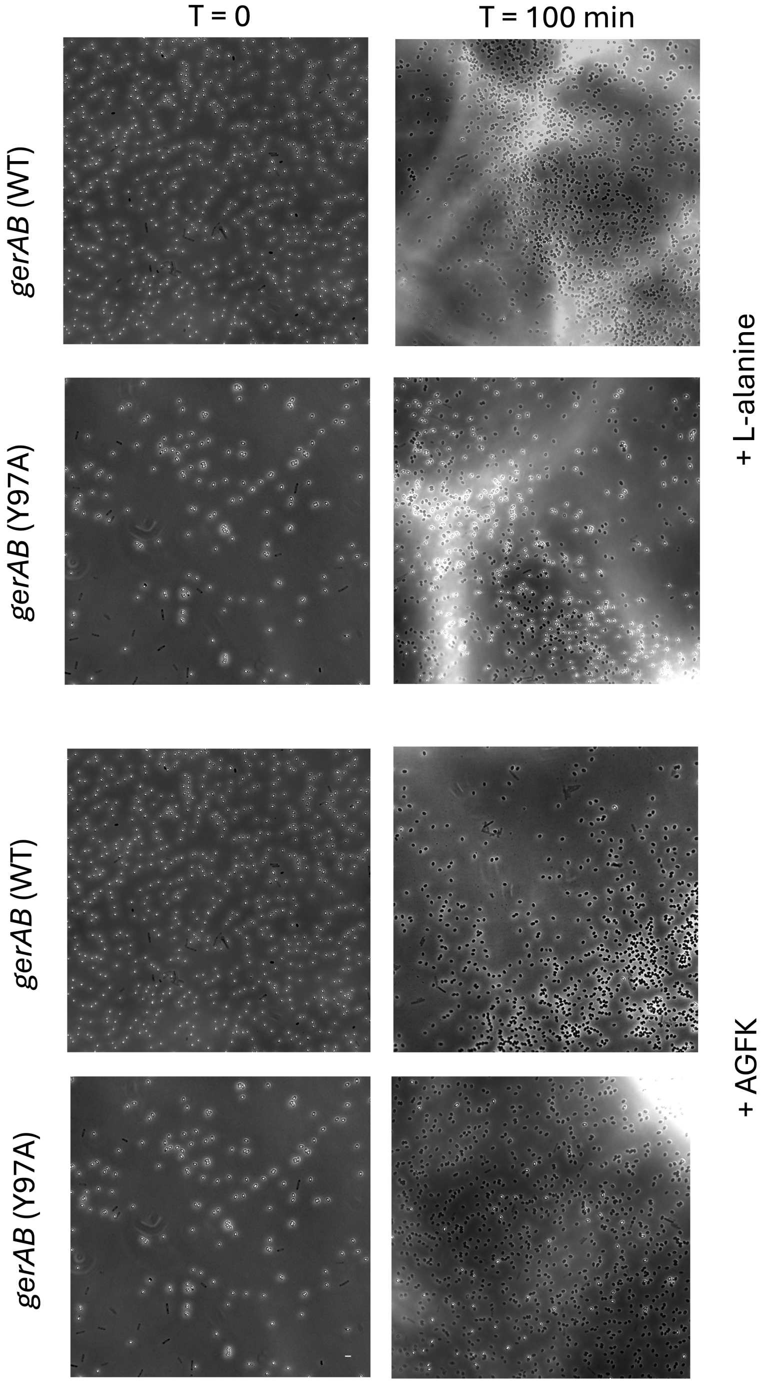


**Figure S1.** Uncropped microscopy image of representative germination assay of wt and Y97A spores before and after 100 min incubation with L-alanine or AGFK, respectively. Scale bar, 2 µm.

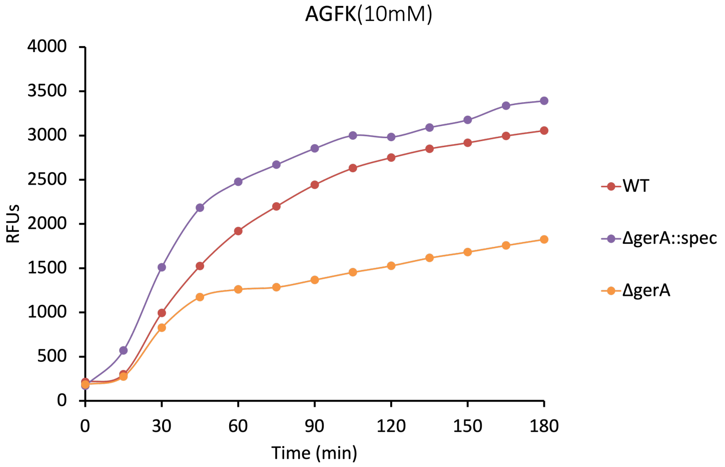


**Figure S2.** Germination of *ΔgerA* strain compared to WT based on DPA release experiment. *ΔgerA* strain showed partial germination compared to WT strain. Since certain point mutation on GerAB has the same effect in AGFK germination than *ΔgerA* strain, it strengthens our observation that the structural integrity of germinosome was affected in strains harboring those point mutations. However, *ΔgerA::spec* strain exhibited higher level of germination compared to WT. Albeit out of the scope of the current study, it seems that spectinomycin resistance marker inserted in genome could affect AGFK germination for some reason.

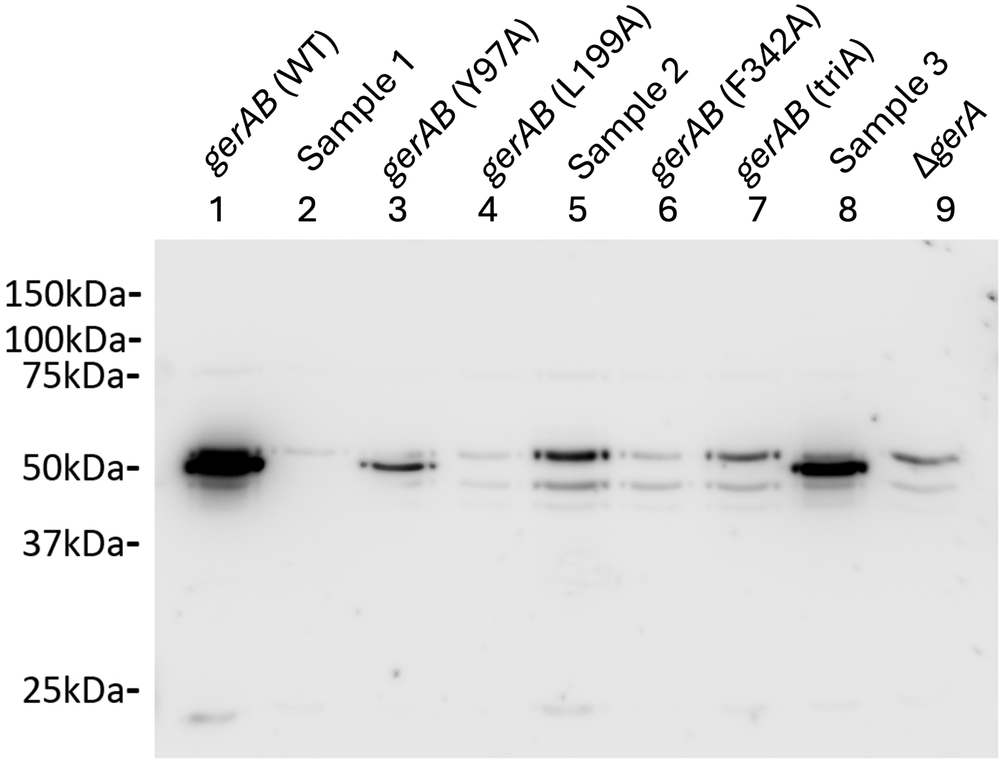


**Figure S3.** Uncropped western blot of *B. subtilis* PY79 wt and mutant spore proteins with GerAA antibody. Sample 1, 2 and 3 were removed from Figure 5 in main text, as they are irrelevant from the current study.
